# Assessing the relationship between cognitive empathy, age, and face perception using a sequential Bayesian analysis

**DOI:** 10.1101/2020.09.11.292706

**Authors:** J.D. Kist, R.A.I. Bethlehem, Barnaby Stonier, Olivier Sluijters, S. K. Crockford, Elke de Jonge, Jan Freyberg, Simon Baron-Cohen, O. E. Parsons

## Abstract

The ability to correctly identify emotions in social stimuli such as faces is proposed to affect one’s level of cognitive empathy. The Fusiform Face Area shows a heightened neural response during the perception of faces relative to objects. We tested whether neural responses to social stimuli were associated with performance in a measure of cognitive empathy, the ‘Reading the Mind in the Eyes’ Task. To quantify face perception, participants were presented with images during a fast serial presentation task which elicited a steady state visual evoked potential, measured using electroencephalography. A Sequential Bayesian Analysis was used to assess if face specific neural responses were associated with either cognitive empathy or age. Data were collected from a participant group of both neurotypical individuals and individuals on the autistic spectrum. We found no significant relationship between the face-specific neural signature, cognitive empathy or age. This study highlights the efficiency of the Sequential Bayesian Analysis as an effective method of participant recruitment.

## Introduction

Autism spectrum condition (ASC) is a highly varied neurodevelopmental condition characterized by differences in sensory perception, difficulties with social interactions and the display of repetitive behaviors and obsessions (American Psychiatric Association 2013). The atypical patterns found in the sensory perception in ASC have been suggested to affect social cognition and consequently behavior in a bottom-up manner (Robertson and Baron-Cohen 2017; Dellapiazza 2018). The ability to engage in successful social interaction relies not only on bottom-up processes such as sensory perception, but also on more internal, higher level processes such as cognitive empathy. Cognitive empathy is defined as the ability to recognize and understand mental states in oneself or other people. Levels of cognitive empathy vary throughout the entire population, along a bell curve, however autistic individuals are consistently found to have lower levels of cognitive empathy than non-autistic individuals (Jones et al. 2010; Rueda, Fernández-Berrocal, and Baron-Cohen 2015; Greenberg, Warrier, and Allison 2018).

The perception of emotion in other people is at the core of cognitive empathy: whether we perceive an emotion correctly in someone else determines how we react to them (Thye et al. 2018; S. Baron-Cohen 2011). A large amount of information concerning an individual’s emotion and identity is contained in the face (Williams and Cross 2018; Simon Baron-Cohen et al. 1997). Because of this, the ability to read facial information quickly is an important, evolutionarily beneficial trait and is vital to facilitating social interactions (Walter et al. 2005); (Kanwisher and Yovel 2006). It has been suggested that the perception of faces includes some partly innate mechanisms and that face saliency during perception occurs early in development (Schultz et al. 2003; Rossion and de Heering 2015).

A network of neuro-anatomical areas is responsible for the visual and emotional perception of faces, with the fusiform face area (FFA) playing a key role. In fMRI studies, the FFA has repeatedly been identified as the most active region during face perception tasks (Pierce et al. 2001). Consequently, the face-specificity hypothesis proposes that there exists a distinct mechanism for face processing (Kanwisher and Yovel 2006; Rhodes et al. 2004). The recognition of faces may therefore not only be functionally distinct from object recognition, but also anatomically segregated (Pitcher, Duchaine, and Walsh 2014).

An alternative theory on the functional specificity of the FFA suggests that activity may be experience-related (Gauthier et al. 2000). The Domain-General theory suggests that the FFA is active in the processing of various stimuli, not just restricted to faces, as long as we can build experience with these objects. Since we continuously build experience with the faces we see around us, the FFA may just be primed to react strongly to faces due to this long-present experience. Functional MRI studies have found evidence suggesting that face-related activity in the FFA is likely to become stronger with age due to a continuous increase of experience with faces (Nordt et al. 2018; Natu et al. 2016).

However, it has also been shown that the perception of faces and the recognition of both identity and emotion decline after a certain age (Ruffman et al. 2008; Mather 2016). Neural networks necessary for face processing also change with age (Mather 2016; Thomas et al. 2008). The superior temporal gyrus (STG), orbitofrontal cortex (OFC) and the amygdala have been suggested as part of the entire emotional brain (Baron-Cohen et al. 1999). Aging of these areas, particularly the STG (including the FFA and the pSTS) may affect the perception of faces during decline. In a diffusion tensor imaging (DTI) study by Thomas et al. it was found that the integrity of white matter tracts in the right FFA reduced with age (Thomas et al. 2008). The decline of neuro-anatomical regions which are involved in the perception and processing of social stimuli from the visual cortex to higher processes in the frontal regions, may therefore contribute to the decline of emotion recognition and self-reported empathy in aging individuals. The effect of age on face perception may therefore be an important factor to address while comparing the two hypotheses.

Perceptual categorization of faces is commonly found from a young age in human infants, and remains present throughout development. In a study by Rossion and de Heering (2015) it was shown that neurotypical infants as young as four months showed heightened neural responses elicited by the perception of faces. While infants from 3 years of age already respond stronger to faces in the FFA, reduced eye-contact and a diminished motivation to engage in social interaction (Pierce et al. 2001; Schultz et al. 2003; Moriuchi, Klin, and Jones 2017; Tanaka and Sung 2016) may result in individuals with ASC being ‘face-inexperienced’.

In order to test whether face-related activity in the FFA is directly related to cognitive empathy and/or age we set up an electroencephalography (EEG) study using a paradigm similar to the one used by Rossion & de Heering (2015). This paradigm involves the periodical presentation of social stimuli within sequences of non-social stimuli and quantifying a face-specific neural response based on the relative evoked potentials of the two stimuli types. We then used an analysis called the Sequential Bayes Factors (SBF) to assess the relation between face-related activity, cognitive empathy and age (Schönbrodt et al. 2015). SBF allows for the sequential computation of Bayes factors for the correlation between two factors until a predefined level of evidence is attained. The SBF approach was used to allow gradual and simultaneous participant-recruitment and analyses, with the option to stop recruitment when the results produced clear converging evidence in support of either the null or alternative hypotheses.

## Results

We collected data from a group of largely neurotypical individuals (3 individuals reported a diagnosis of ASC) as part of a larger ongoing EEG study. Participants (N = 56) completed the Autism Spectrum Quotient (AQ) and the ‘Reading the Mind in the Eyes’ Task. We measured the face specific neural signature using EEG. Participants’ ages ranged from 17 to 66 years old (26 females and 30 males, mean age = 27.5 sd = 12.3).

### EEG Set-up and Task

We recorded the EEG data using a BioSemi active two 64-electrode system. Participants wore a biosemi cap with a 20-10 layout system, in which electrodes were inserted with saline-gel filled plastic electrode holders. We used a fast serial visual presentation (FSVP), as used in the study by Rossion and de Heering (2015, 9–10). The task exposes the viewer to a series of images at a steady frequency (see methods and figure 8.A. (Rossion and de Heering 2015) for details). The images used portray neutral scenes, such as objects and houses. These were presented at a 6 Hz frequency, fading in and out using a sinusoidal contrast modulation. The computer screen that we used during the task presentation has a 144 Hz refresh rate in order to facilitate the rapid contrast modulation.

**Figure 1.**
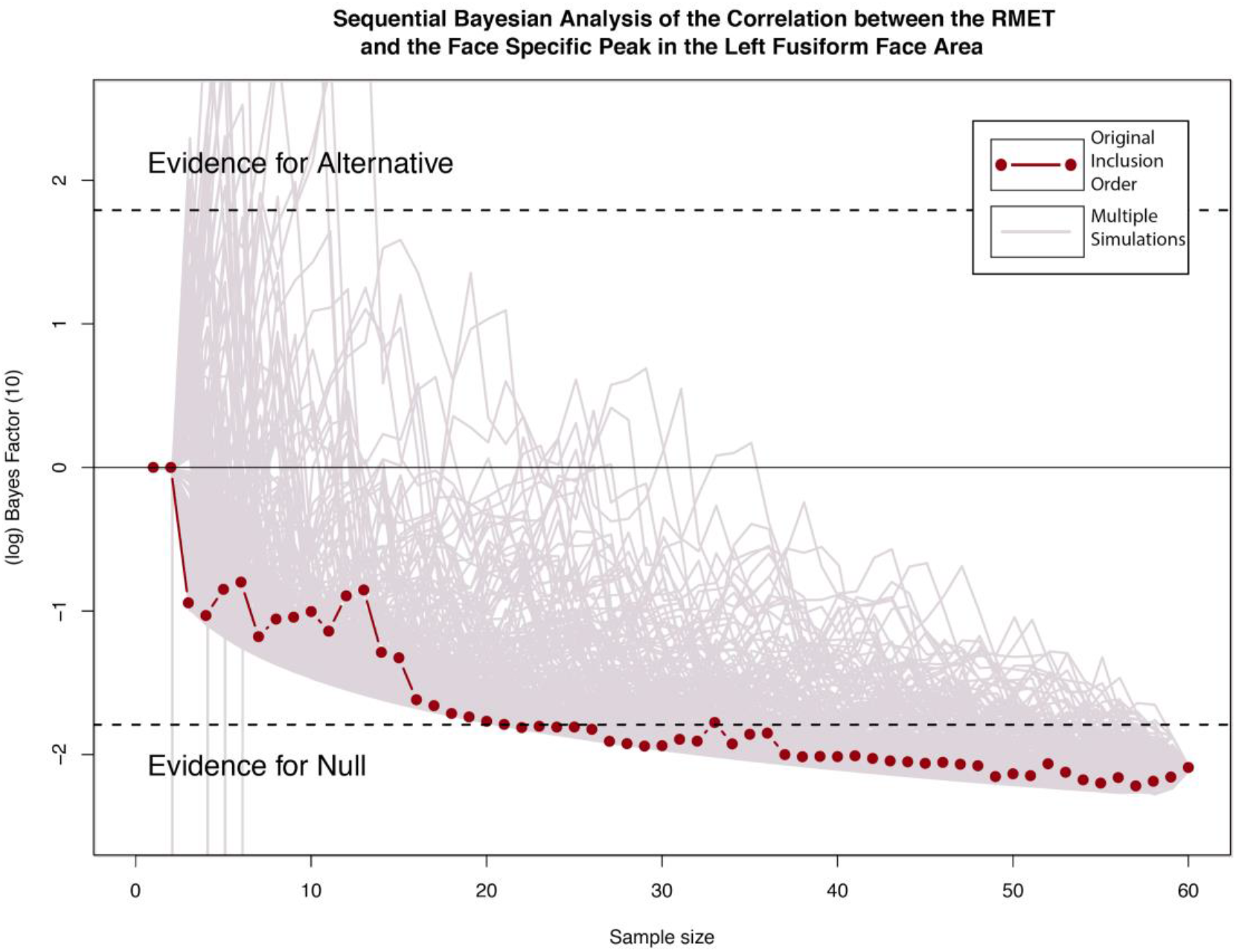
Sequential Bayesian Analysis of the Correlation between the RMET and the Face Specific Peak in the Left Fusiform Face Area. Accumulation of evidence throughout data collection for the correlation between RMET scores and the Face Specific Peak in the Left FFA (Final Bayes Factor: *BF_10_*= 8.084, *BF_01_*= 0.1234, *Log_10_*(*BF_10_*) = −2.090) The convergence of the multiple simulations indicates that there exists decisive evidence in favour of the null hypothesis.

**Figure 2.**
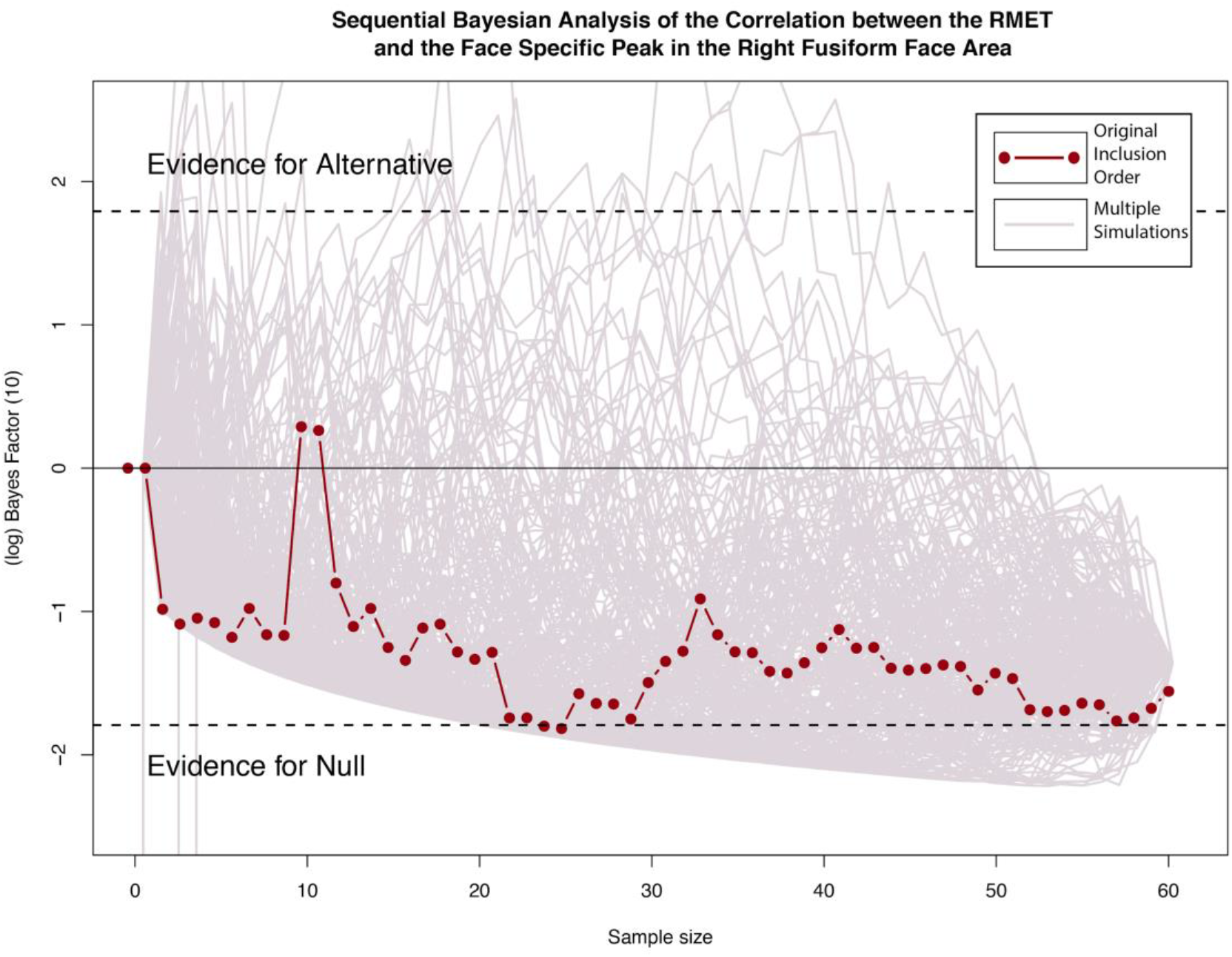
Sequential Bayesian Analysis of the Correlation between the RMEt and the Face Specific Peak in the Right Fusiform Face Area. Accumulation of evidence throughout data collection for the correlation between RMET scores and the Face Specific Peak in the Right FFA (Final Bayes Factor: *BF_10_*= 3.9024, *BF_01_*= 0.256, *Log_10_*(*BF_10_*) = - 1.362) The convergence of the multiple simulations indicates that there exists strong evidence in favour of the null hypothesis.

**Figure 3.**
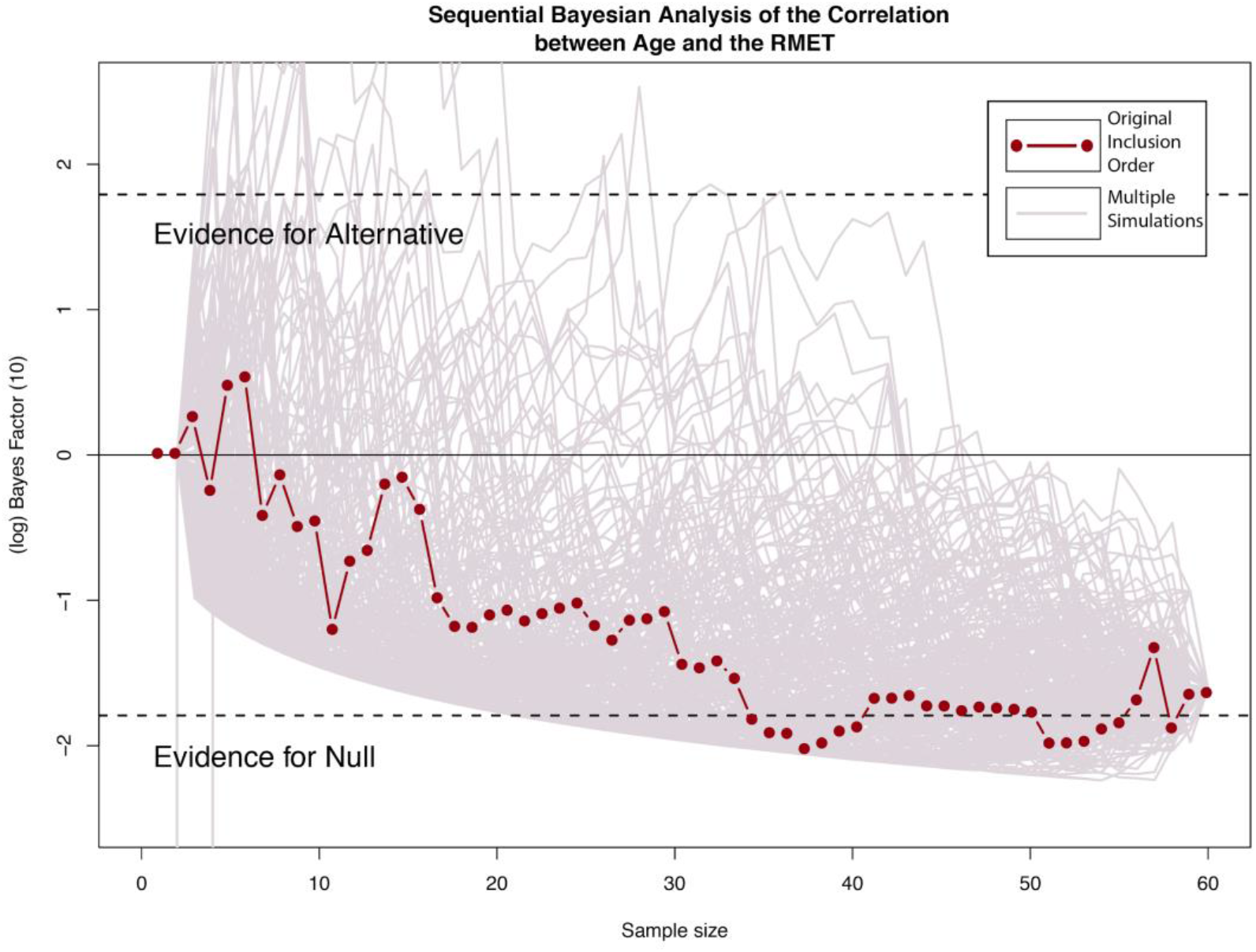
Sequential Bayesian Analysis of the Correlation between Age and the RMET. Accumulation of evidence throughout data collection for the correlation between Age and the RMET scores (Final Bayes Factor: *BF_10_*= 4.970, *BF_01_*= 0.2012174, *Log_10_*(*BF_10_*) = −1.603). The convergence of the multiple simulations indicates that there exists strong evidence in favour of the null hypothesis.

**Figure 4.**
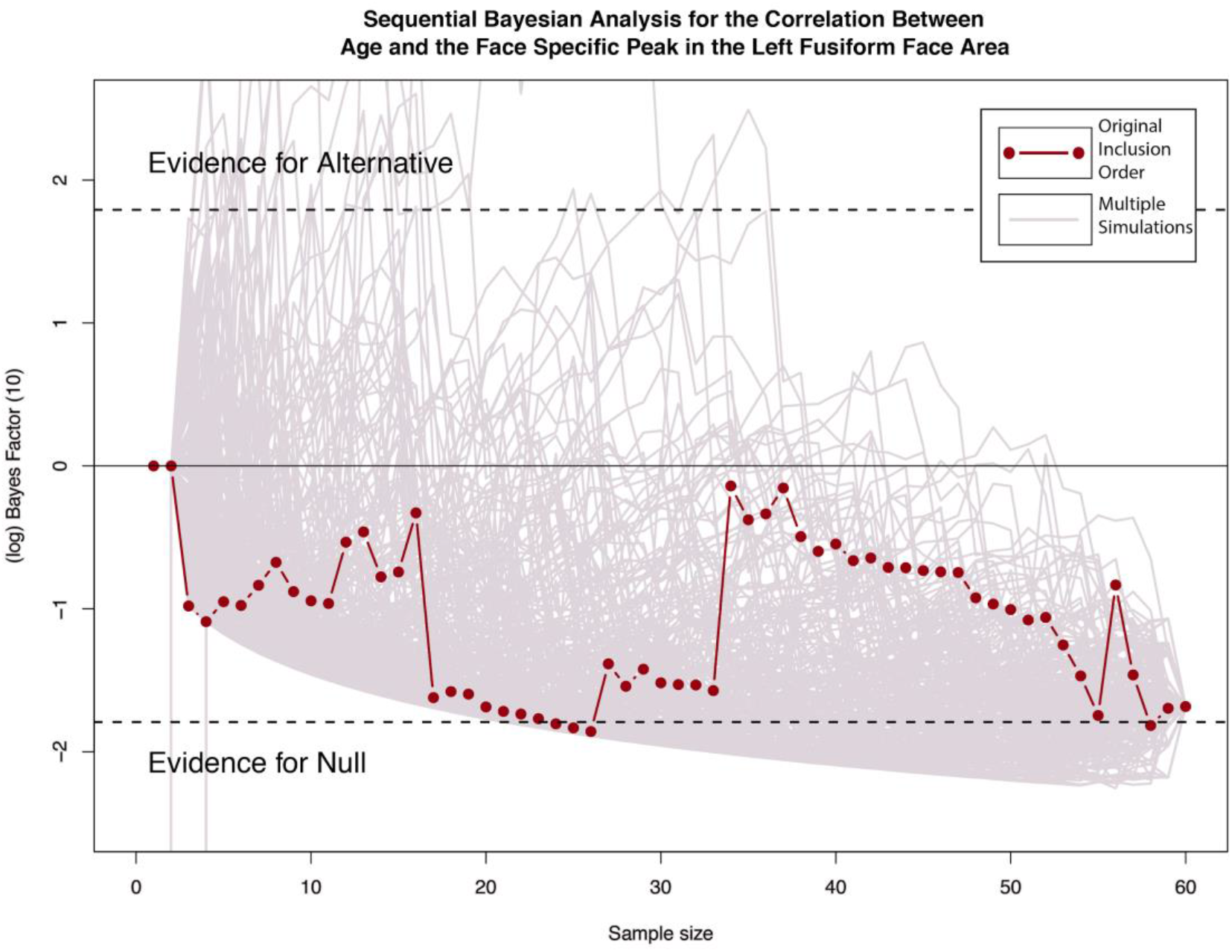
Sequential Bayesian Analysis for the Correlation Between Age and the Face Specific Peak in the Left Fusiform Face Area. Accumulation of evidence throughout data collection for the correlation between Age and the Face Specific Peak in the Left FFA ((*BF_10_*=5.390, *BF_01_*= 0.1856731, *Log_10_*(*BF_10_*) =−1.684). The convergence of the multiple simulations indicates that there exists strong evidence in favour of the null hypothesis.

**Figure 5.**
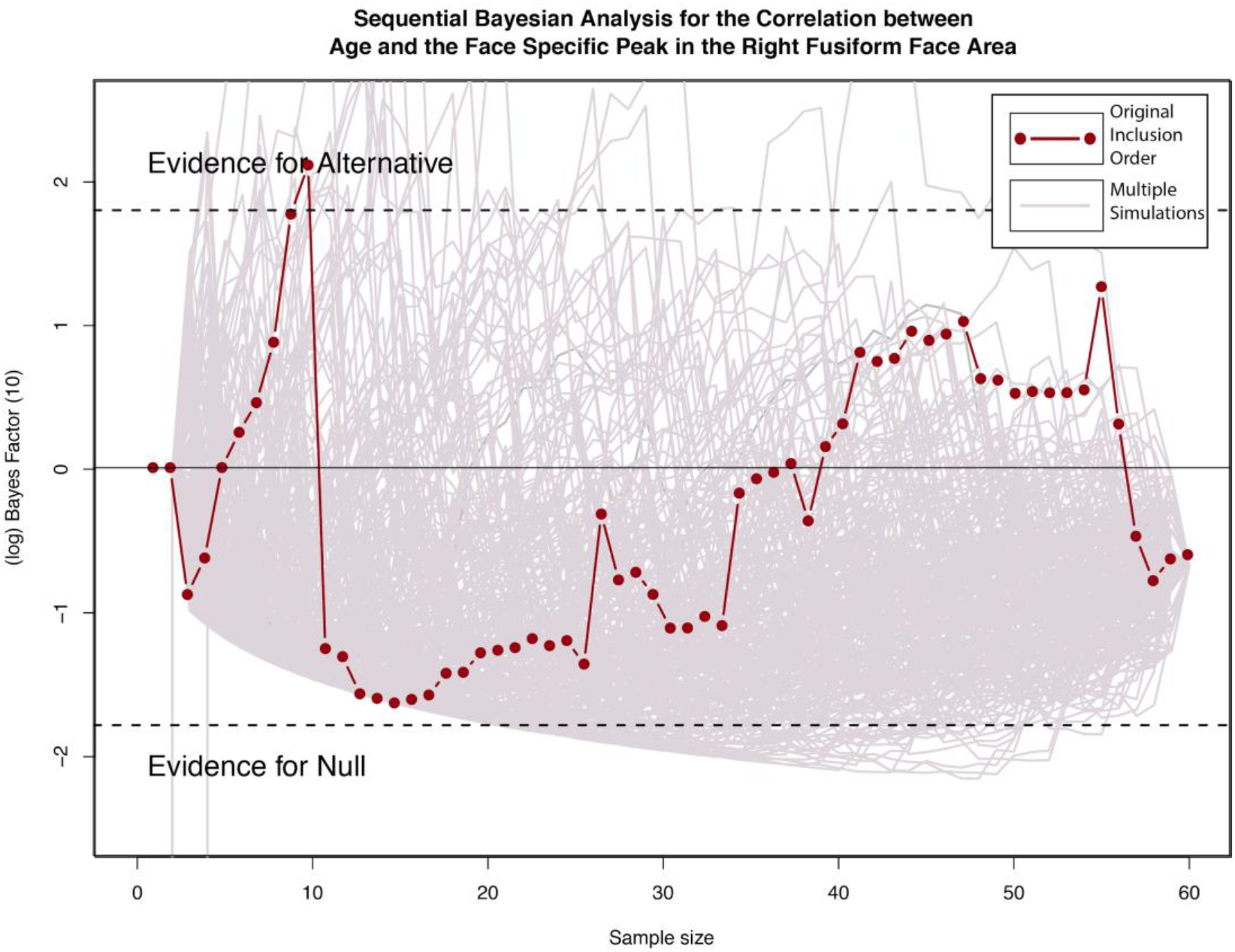
Sequential Bayesian Analysis for the Correlation Between Age and the Face Specific Peak in the Right Fusiform Face Area. Accumulation of evidence throughout data collection for the correlation between Age and the Face Specific Peak in the Right FFA ((*BF_10_*=1.954, *BF_01_*= 0.5118, *Log_10_*(*BF_10_*) = −0.670) The convergence of the multiple simulations indicates that there exists weak evidence in favour of the null hypothesis.

**Figure 6.**
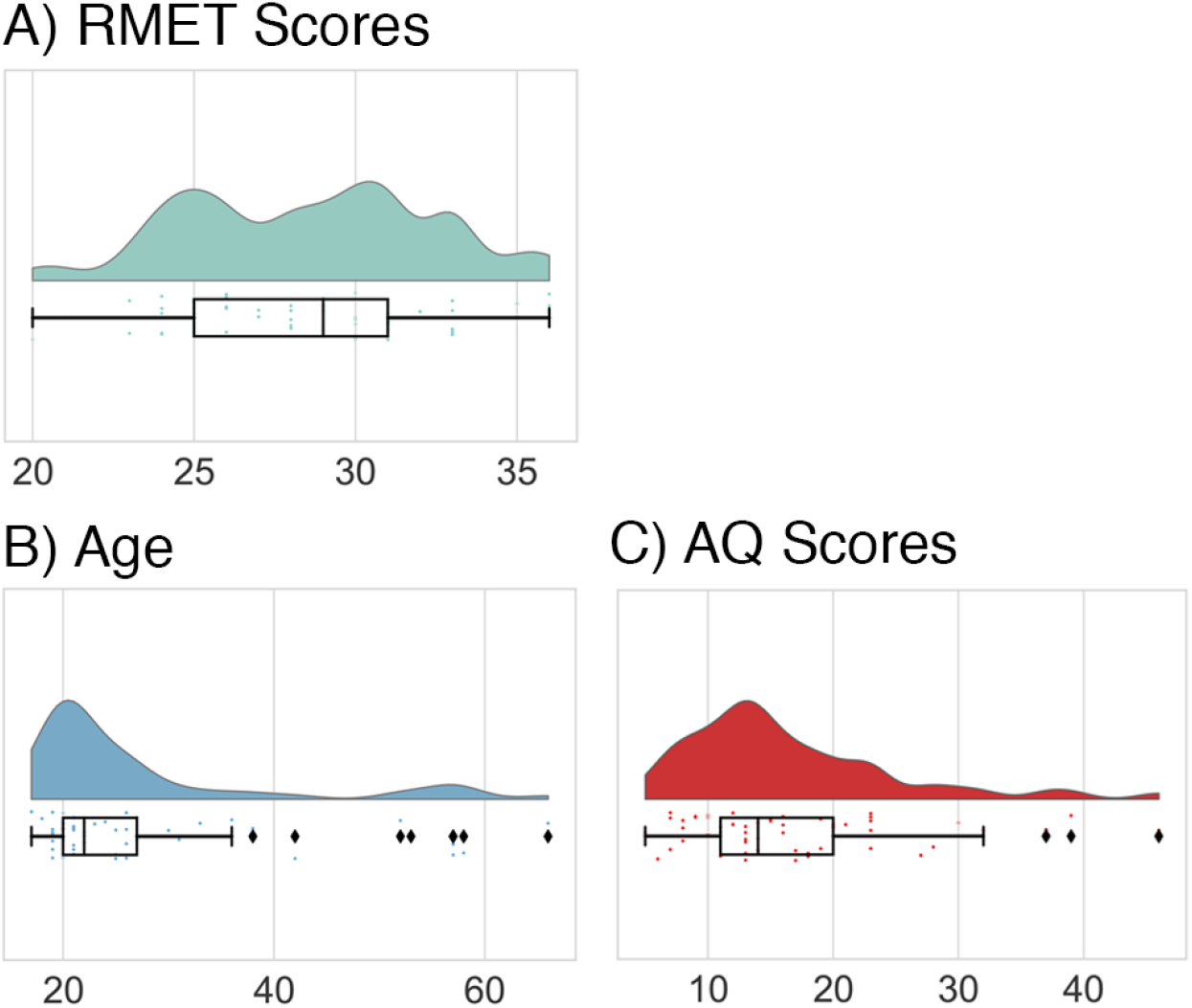
Distribution of RMET, AQ and Ages of the included participant group (n = 56) A) RMET scores B) Age C) Autism Quotient Scores

**Figure 7.**
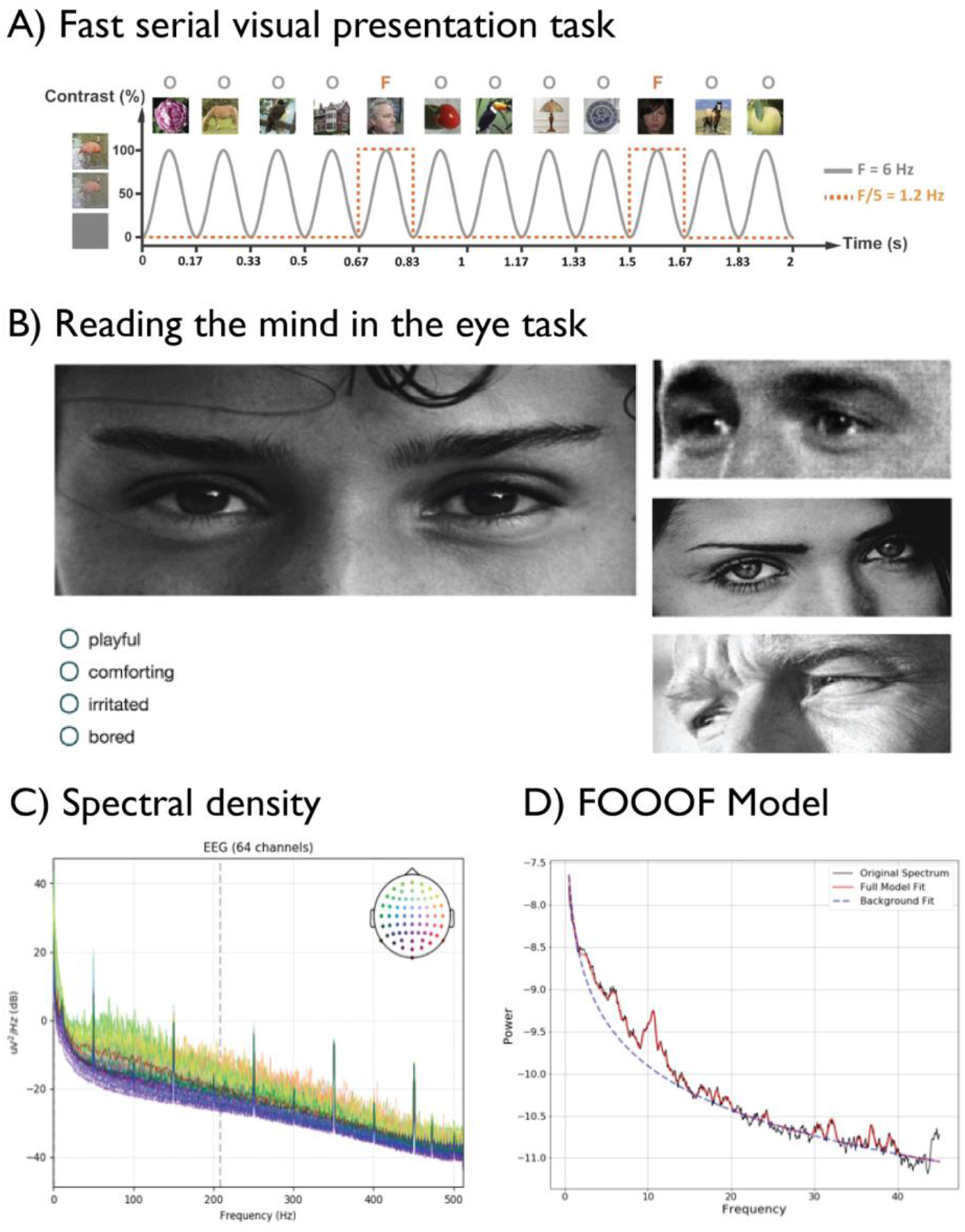
A Fast Serial Visual Presentation Task. B. The Reading the Mind in the Eyes Task. C. Spectral Density. D. FOOOF Model.

**Figure 8.**
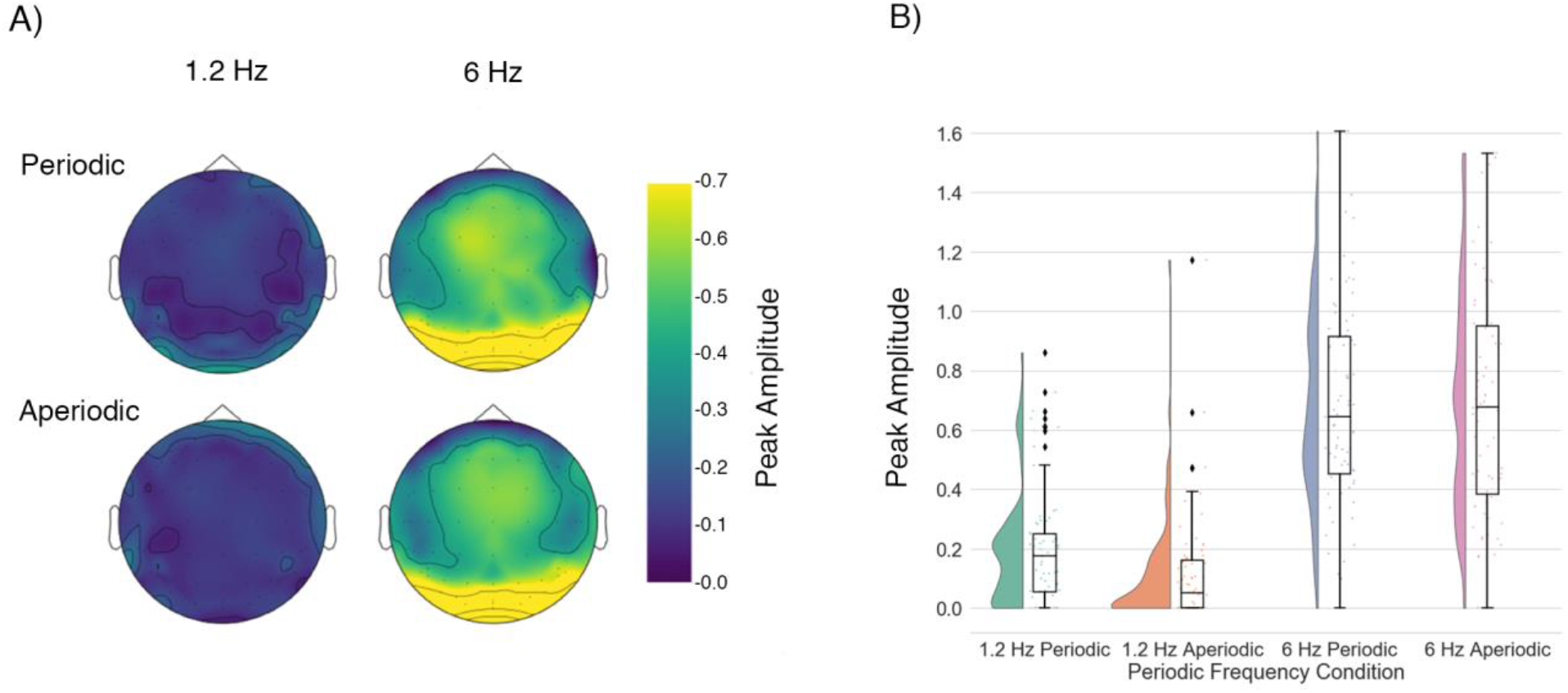
**A)** Topo Maps of the peak amplitude per periodicity and frequency. **B)** RainCloud Distribution of the average peak amplitude per periodicity and frequency in the right FFA periodic baseline peak (M: 0.742659, SD: 0.331077), aperiodic baseline peak (M: 0.551727, SD: 0.325041) periodic face-specific peak (M: 0.207213, SD: 0.205435) aperiodic face-specific peak (M: 0.118524, SD 0.192918)

The FSVP task consists of 16 trials including 8 periodic trials, in which every fifth image (at 1.2 Hz) displays a face image instead of a neutral image. Furthermore, there were 8 aperiodic trials in which face images were presented at random intervals. Each trial lasted 14 seconds. With the use of an FSVP approach, we can isolate the exact frequency at which stimulation is produced to measure the elicited response. As such it allows for both a predictive and an objective analysis.

The periodic exposure to the stimulus allows the isolation of a specific voltage peak of the signal at the determined frequency, known as: a steady state visual evoked potential (SSVEP). We isolated the SSVEP and used it for the computation of the face-specific neural signature for further analyses. Furthermore, we downsampled the signal to 1024Hz during recording, as this provided enough temporal resolution to detect the SSVEP.

### The Sequential Bayesian Plots

We plotted the association and sequential bayes of the separate relationships between the face specific neural activity, cognitive empathy and age to assess the sampling effects, and to evaluate the consistency across simulations.

### The Relationship Between Cognitive Empathy and Face Specific Neural Activity

Firstly, we plotted the association and sequential bayes of the relationship between the face specific neural signature and cognitive empathy, as measured by the RMET. We found decisive evidence for a lack of relationship between the face specific peak in the left fusiform face area and RMET scores (Final Bayes Factor: 8.084) (Figure 1.).

We also found strong evidence for the lack of relationship between the face specific peak in the right fusiform face area and RMET scores (Final Bayes Factor: 3.9024) (Figure 2.), meaning there was no significant correlation between cognitive empathy and facial perception as found in the fusiform face area in both hemispheres. As can be seen in both Figures 1 and 2 the multiple simulations indicate that there was a high chance of false positives in lower sample sizes. This indicates that incorrect conclusions regarding the existence of a relationship could easily be drawn if an insufficient number of participants were included in the analysis.

### The Relationship Between Cognitive Empathy and Age

Secondly, we plotted the association and sequential bayes of the relationship between age and cognitive empathy. In this dataset there seems to be no significant relationship between age and cognitive empathy. We found strong evidence for a lack of relationship (Final Bayes Factor = 4.970), indicating that age may not influence cognitive empathy (Figure 3.). Again, a substantial number of false positives could be found in the simulations, at a lower number of sample sizes.

### The Relationship Between Age and Face-Specific Neural Activity

Lastly, we plotted the association and sequential Bayes of the relationship between the face specific neural activity and age. We found evidence for a lack of relationship between age and face-specific activity in both the fusiform face areas. Bayes Factors indicate no correlation between age and face specific peak frequency in neither the left (final BF = 5.390, figure 4) nor the right (final BF = 1.954, figure 5) FFA. This indicates that age may not influence the activity in either the left or the right fusiform face area.

In all of the plotted simulations a high number of false positives could be found, although mainly at lower sample sizes. The use of these plotted simulations allows for an unbiased assessment of sampling effects.

## Discussion

The present study aimed to test if there is a correlation between a face-specific neural signature and cognitive empathy in a typical participant group. With the use of a validated behavioural measure of cognitive empathy, the ‘Reading the Mind in the Eyes’ Test, a significant relation between the two could have established the face-specific neural signature as a potential biological neuromarker of cognitive empathy (Simon Baron-Cohen et al. 2001). Our Sequential Bayesian Analysis indicated that with our participant group there was strong evidence against such an association.

There may be several reasons for the lack of relationship between the RMET and the face-specific neural signature in this dataset. Firstly, the face-specific neural signature used in this study, the steady state visually evoked potential during face presentation in the FFA, is a low level aspect of visual perception of faces (Rossion and Boremanse 2011). Therefore, face-specific neural signature may be too low-level a process to have an immediate relationship with a higher-level concept such as that of cognitive empathy.

It is possible that at this low-level stage of perception no deviations exist, but that instead an altered integration of perceived stimuli further along the visual processing stream influences cognitive empathy. The current FSVP task concerns purely the classification of faces versus neutral images. No emotionally salient aspects or features were perceived that required higher levels of integration, such as emotional recognition, gaze following or holistic face-processing. While there may be individual differences at the low-level perception of faces vs non-faces, these may not have a direct relationship with the level of cognitive empathy. This explanation of our results aligns with the Weak Central Coherence theory (Happé 1996; López et al. 2004), which posits that fundamental perception is intact or even heightened for simple features in individuals with ASC, yet that the integration of these perceptions at a higher processing level, for example during emotion processing, is affected. As such, there may be no altered sensory perception affecting cognitive empathy, but an altered integration of sensory input. As the RMET requires higher-level processing (Fernández-Abascal et al. 2013; Preston et al. 2007; Moor et al. 2012), it is likely that the local processing bias has no direct link with an individual’s performance on this task, but rather the inability to integrate the separately perceived social stimuli.

Furthermore, we studied a participant group of mostly neurotypical individuals. While we have approached cognitive empathy as a spectrum, in which no real cutoff for dysfunction of cognitive empathy was set, it may be the case that a relationship between facial perception and cognitive empathy is only noticeable at pathological levels. The lack of individuals with an actual ASC diagnosis in our participant sample, may have prevented us from finding a significant relationship between face perception and cognitive empathy, as cognitive empathy has been found to be significantly lowered in individuals with ASC (Peñuelas-Calvo et al. 2019; Schultz et al. 2003; Schultz 2005). Therefore we suggest that future studies may focus their attention on the possible relationship between holistic face-processing and cognitive empathy, and include a higher number of participants with an ASC diagnosis.

According to the experience-hypothesis, one would expect a better ability to judge emotion and facial expression with older age. Aging has been proven to lead to an increase in cognitive empathy up to a certain age (Thomas et al. 2008; Ruffman et al. 2008; Nordt et al. 2018); with older age cognitive decline and thereby a decrease of cognitive empathy is common, as neural capacity and executive functioning decrease. In our sample we found evidence for the lack of a relationship between age and cognitive empathy, as well as age and the face-specific neural signature. The evidence for the lack of a relationship seems to contradict the experience hypothesis of the fusiform face area. The distribution of age in the current sample, however, may have prevented us from finding this relationship (Figure 6.B.). Since the distribution of age in our sample is skewed, this evidence may be futile. Future studies assessing this hypothesis therefore need to consider age distribution when assessing the effect of age on neural signatures and cognitive empathy.

This study was designed around the premise that steady state visual evoked potentials (SSVEP) could be triggered in the FFA, through a fast serial visual presentation (FSVP) of imagery, as was similarly done in a previous study conducted by Rossion et al. (2015). The detected SSVEP indicated that there is a stronger response in the FFA towards faces rather than objects (Figure 8.B.). This result suggests that the methodology is valid, and that the FFA shows a heightened response to face-imagery, as has been found in previous studies (Kanwisher and Yovel 2006; Rossion and Boremanse 2011). However, while this finding is in line with the face-specific hypothesis, it is not sufficient to disprove the experience-related hypothesis (Gauthier et al. 2000; Rhodes et al. 2004).

Therefore we have shown the functionality of an EEG paradigm based on the SSVEP, the efficiency of the analysis, and its ability to eliminate possible participant recruitment order effects. The sequential Bayesian analysis has facilitated analysis during the on-going recruitment of participants, by allowing us to bypass the process of setting a predefined inclusion number of participants before analysis could take place. Thus, the use of an SBF can allow for flexibility in sample-planning and facilitate this process when participants are harder to recruit, for example in rare clinical samples. In the case of unclarity in terms of existing relationships, it is possible to include more data-points in an unbiased manner while increasing the accuracy of the effect size (Schönbrodt et al. 2015). Furthermore, the SBF makes for an intuitive tool to account for sampling order. Especially in lower sample sizes the sequential Bayesian simulation showed that there are likely quite a few false positives in our sample. This could be a result of order-effects in smaller sample sizes. Thus, without the SBF analysis it would be possible to make premature conclusions on existing relationships, whereas with the use of the SBF we could ensure that there was no evidentiary support for the relationships in question.

In summary, by using sequential Bayesian analysis we found no evidence for an association between cognitive empathy, facial visual perception or age. However, we have highlighted the usefulness of SBF to determine appropriate sample size and stopping criteria. We acknowledge that an association may exist but may be precluded by the current sample distribution. Future studies should consider adopting a similar bayesian framework for robust hypothesis evaluation.

## Materials and Methods

### Data Distribution

We visualized the distribution of data points within our sample using Raincloud Plots. We created these with the use of code developed by Micah Allen (Allen et al. 2019). While the RMET scores (Figure 6.A) show a rather normal distribution, the Age and AQ scores (Figure 6.B and 6.C respectively) show a fairly skewed data point distribution. Further information on the correlation between these measures can be found in the Supplementary Materials.

### The Reading the Mind in the Eyes Task

We measured cognitive empathy by means of the Reading the Mind in the Eyes Test (RMET) (See Figure 7.B). This test has been validated as an appropriate behavioural measure of cognitive empathy (Fernández-Abascal et al. 2013). It measures the extent to which individuals can attribute and recognise people’s facial expressions solely via a cropped facial area around the eye (Simon Baron-Cohen et al. 2001). The RMET has been found to have good reliability, as individuals with ASC consistently score lower on the task (Fernández-Abascal et al. 2013; Baron-Cohen et al. 1997). As such the RMET has often been used in various types of research studying the mechanisms behind cognitive empathy (Warrier et al. 2018; Baron-Cohen et al. 2015; Greenberg et al. 2018).

### Processing of the EEG signal

We preprocessed and analysed each acquired EEG dataset using MNE software in Python 3.6 (available at https://mne.tools/dev/install/index.html and https://www.python.org/downloads/release/python-366/) (Gramfort et al. 2013). The raw data was separated in 14 second epochs which were assessed for quality. Epoch quality was assessed both manually and using AutoReject: Bad channels were interpolated where possible, using the *Random Sample Consensus (Ransac)* function of AutoReject (Jas et al. 2017; Bigdely-Shamlo et al. 2015). The Ransac algorithm determines the data quality of each channel and, in case of bad quality, repairs these. Thereafter, we determined the quality of each epoch-object and cleaned the dataset of bad epochs. If less than 11 out of 16 epochs remain from a dataset (68.75%), we removed it from analysis. At this stage 8 datasets were excluded due to noisy epochs. The raw data was filtered (0.5 - 15), demeaned and underwent linear detrending. We performed a Fast Fourier Transform (FFT) to construct a data structure for the SSVEP analysis, determining the signal to noise (SNR) ratio during the frequencies of our interest (6Hz and 1.2Hz). Jan Freyberg developed the code that we used in order to do this (https://github.com/janfreyberg/face-perception-ssvep) We computed the SNR by comparing the signal level during the stimulus to the signal level without the stimulus.

#### Controlling for an Aperiodic Signal

EEG signals are inherently noisy, and the amplitude of this noise changes with frequency. A large component of this signal is proportional to the inverse of the frequency (1/f), and is therefore often called the aperiodic component of neural power spectra. Given that there are individual differences in the parameters of this noise (see Voytek et al, 2015), we aimed to subtract this noise signal from each participant’s EEG spectrum. We used the algorithm Fit Oscillations Of One Over F (FOOOF, (Haller et al. 2018). The aperiodic fit is estimated using linear regression in log-space. This estimate is then subtracted from the power spectrum. The algorithm then iteratively fits gaussians to peaks in the remaining power spectrum until a threshold is reached (Figure 7.D.).

Visualization and Comparison of the face-specific and baseline neural signature We used peaks identified by the FOOOF algorithm as estimates of neural oscillation at the stimulation frequencies (1.2 and 6 Hz). In Figure 8.A. visualization of the average peak amplitude over participants per condition, is plotted as a topo map on the scalp. Localization of the neural activity during the frequencies of interest shows increased high activity in the occipital cortex during 6 Hz, activity was similar during periodic and aperiodic trials. In the 1.2 Hz frequency overall activity was lower. During Periodic trials increased activity could be seen in the occipital cortex and in the left and right FFA, as well as in the T8 and TP8 electrodes; the posterior superior temporal sulcus of the right hemisphere.

The distribution of the average peak amplitude in the right FFA for each participant is visualized per condition in figure 8.B. using a Rain-cloud plot method (as translated to Python https://github.com/pog87/PtitPrince) (Allen et al. 2019). An independent t-test of the baseline peak amplitudes over periodic (M: 0.74, SD: 0.33) and aperiodic trials (M: 0.55, SD: 0.32) differed significantly (t =4.00, p=.0001). The face-related peak amplitudes have a lower peak in general due to the lower presentation frequency. The plot shows that the face-specific periodic peak distribution (M: 0.21, SD: 0.21) is significantly higher (t=3.01, p=0.004) than the aperiodic peak distribution (M: 0.12, SD 0.19). Therefore, looking at the peak amplitude of the periodic face-frequency, it can be concluded that there is a clear SSVEP in response to face-images as compared to neutral images.

#### Sequential Bayesian Factor

The evidence for or against the null hypothesis is determined through the calculation of the Bayesian Factor (BF). The BF is the ratio of one hypothesis being more likely than the other. The BF10 is the ratio of the likelihood of the alternative hypothesis (H1) versus the likelihood of the null hypothesis (H0). We chose a threshold of | log(BF10) | > 1.8, reflecting either hypothesis being 63x higher than the other. According to Kass and Raftery (1995) this indicates ‘strong’ evidence. Sequential bayes factor methods allow assessment of the accumulation of evidence using random sampling. Here, instead of simply evaluating the Bayes Factor with each new participant, after every new data point, we simulated the experiment with a new, random order of participants. This facilitates the visualisation of the trend in evidence over the course of the experiment without confounding order effects.

## Contributions

**J.D. Kist:**

Conceptualization
Methodology
Software
Formal analysis
Investigation
Data curation
Writing
Visualization

**R.A.I. Bethlehem:**

Conceptualization
Methodology
Validation
Investigation
Resources
Data curation
Writing
Supervision
Project administration
Funding

**Barnaby Stonier, Olivier Sluijters, Sarah Crockford, Elke de Jonge:**

Investigation

**Jan Freyberg:**

Conceptualization
Methodology
Software
Validation
Investigation
Resources
Writing (review)
Visualization
Supervision
Project administration

**Simon Baron-Cohen:**

Conceptualization
Resources
Funding
Supervision

**O.E. Parsons:**

Conceptualization
Methodology
Software
Validation
Investigation
Resources
Writing (review)
Visualization
Supervision
Project administration

## Supplementary Materials

An exploratory correlation between the face-specific neural signature in the left and right fusiform face area, the Autism Spectrum Quotient, the RMET scores and age was performed. In Figure 9.A we performed a Pearson Correlation with all the different variables, which can be seen in the heatmap. In Figure 9.B a heatmap can be seen in which a Spearman Correlation was performed.

**Figure 9.**
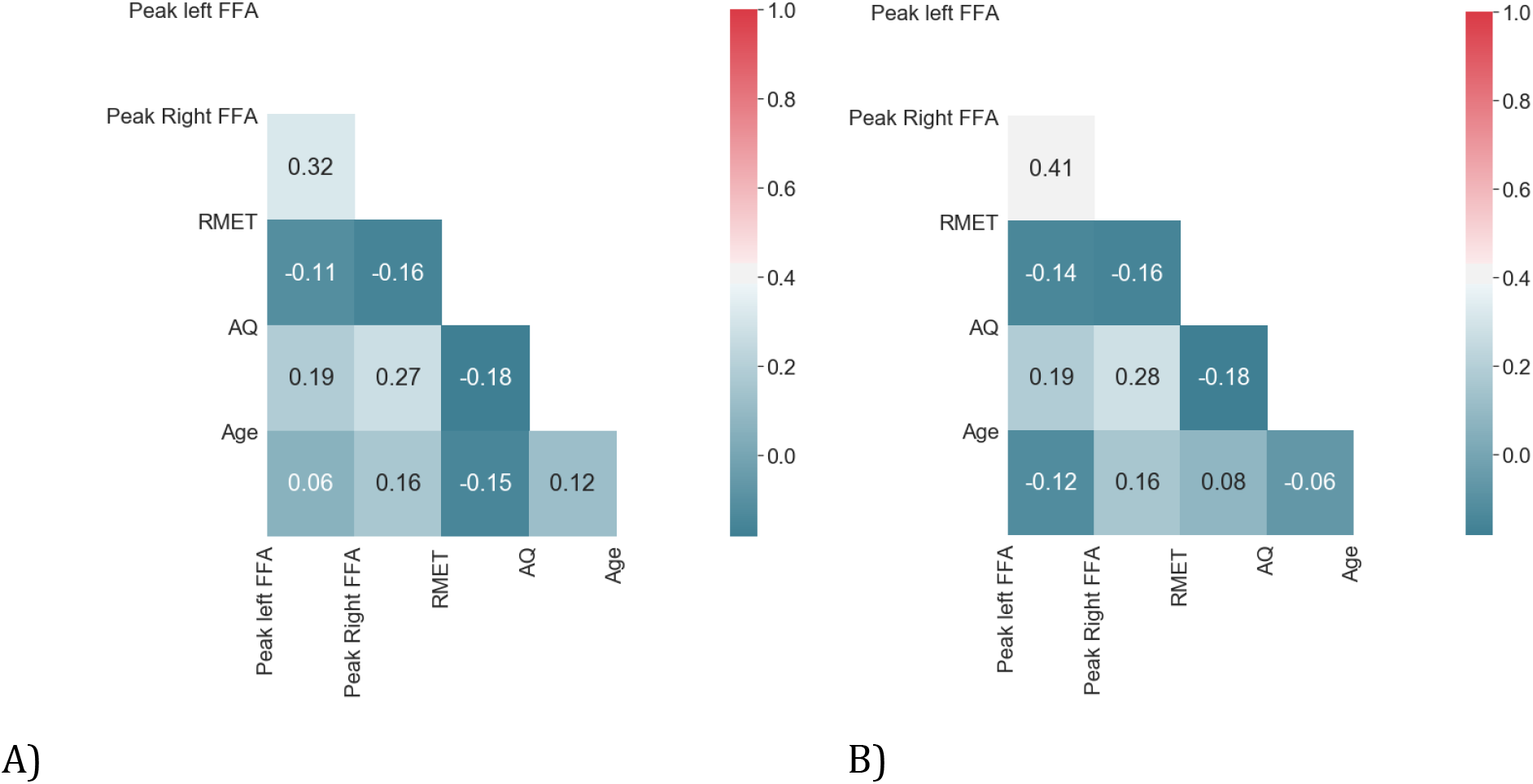
**A)** Pearson Correlation Heatmap. **B)** Spearman Correlation Heatmap (N = 56).

We visualized the different correlations by means of a scatter plot. The different relationships can be seen in Figure 10.

**Figure 10.**
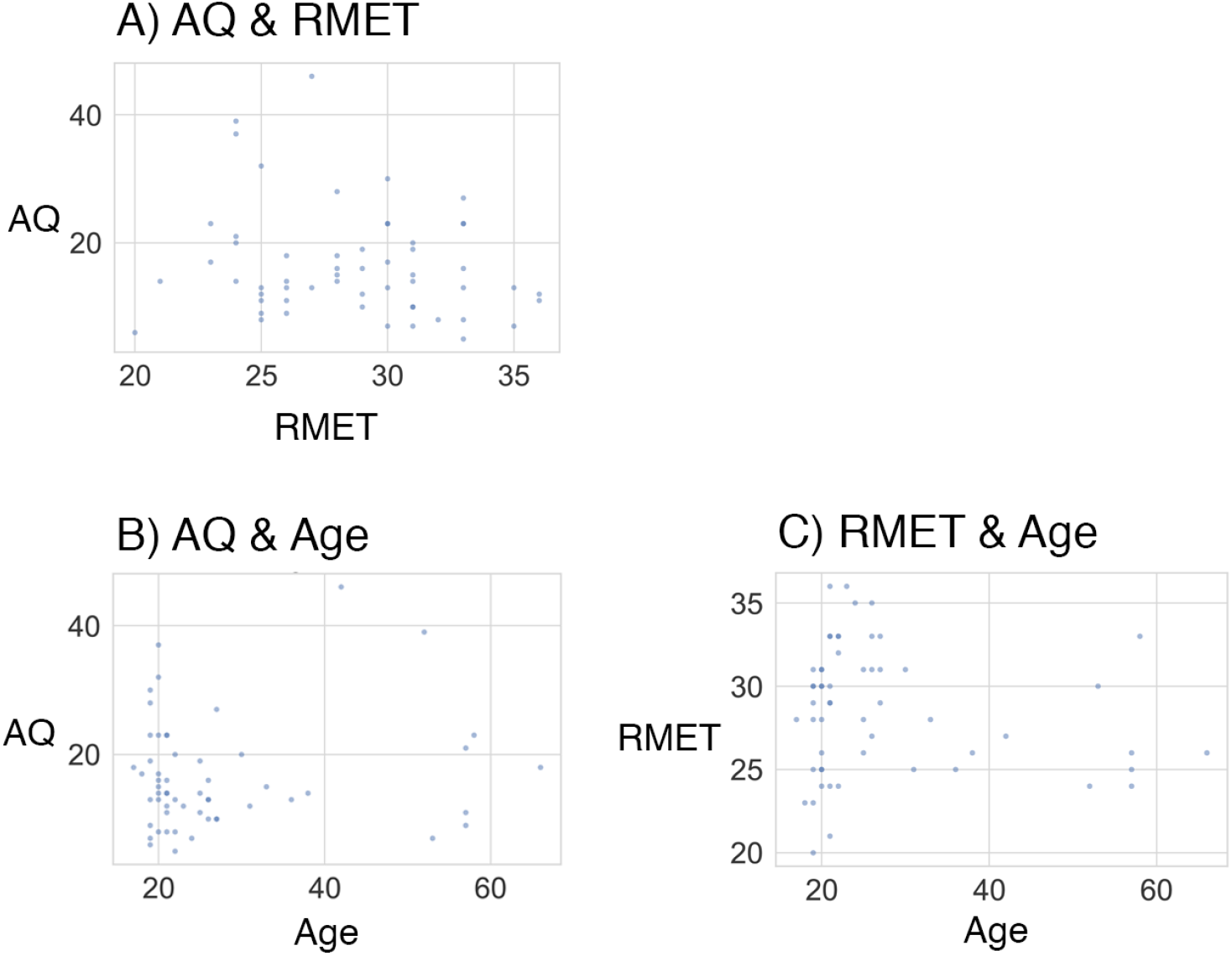
Correlation Scatter Plots. A) Autism Quotient and RMET scores B) Autism Quotient scores and Age C) RMET scores and Age

Furthermore, we performed an analysis to find the electrodes in the EEG set-up that measured the highest ratio of the peak amplitude.

**Table 1.** Electrodes with highest ratio of peak amplitude ratio over participants during periodic trials. Noisy electrodes were excluded if they had a R_squared fit of < 0.9.

|  |  |  |  |  |  |
| --- | --- | --- | --- | --- | --- |
| 1 | FP2 | ventromedial PreFrontal Cortex (vmPFC) | 6 | FP1 | ventromedial PreFrontal Cortex (vmPFC) |
| 2 | AF8 | Right Dorsolateral Prefrontal Cortex | 7 | FT8 | Orbitofrontal Cortex |
| **3** | AF7 | Left Dorsolateral Prefrontal Cortex | **8** | F7 | Left Inferior Frontal |
| **4** | T8 | Right posterior superior temporal sulcus | **9** | TP7 | Left posterior Superior Temporal Sulcus |
| **5** | TP8 | Right posterior Superior Temporal Sulcus | **10** | P9 | Left Fusiform Face Area |

